# Neural network embedding of functional microconnectome

**DOI:** 10.1101/2021.10.19.464982

**Authors:** Arata Shirakami, Takeshi Hase, Yuki Yamaguchi, Masanori Shimono

## Abstract

Our brain works as a complex network system. Experiential knowledge seems to be coded into the organism’s network architecture rather than retaining only properties of individual neurons.

In order to be better able to consider the high complexity of this network architecture, extracting simple rules through both automated as well as interpretable analysis of topological patterns will be necessary in order to allow more useful observations of interrelationships within the complex neural architecture.

By combining these two types of analysis methods, we could automatically compress and naturally interpret topological patterns of functional connectivities, which produced electrical activities from many neurons simultaneously from acute slices of mice brain for 2.5 hours [Kajiwara et al. 2021].

As the first type of analysis, this study trained an artificial neural network system called Neural Network Embedding (NNE), and automatically compressed the functional connectivities into only small (25%) dimensions.

As the second type of analysis, we widely compared the compressed features with ~15 representative network metrics, having clear interpretations, including > 5 centrality-type metrics and newly developed network metrics, that quantify degrees or ratio of hubs distanced by several-nodes from initially focused hubs.

As the result, although we could give interpretations for only 55-60% of the extracted features, these new metrics, together with the commonly utilized network metrics, enabled interpretations for 80-100% features, using automated analysis.

The result demonstrates not only the fact that the NNE method surpasses limitations of commonly used human-made variables, but also the possibility that acknowledgement of our own limitations drives us to extend interpretable possibilities by developing new analytic methodologies.

## 1. Introduction

### 1-1. Network architecture of neuronal circuits

Our brain enables us to perform various functions based on various activation patterns realized on complex structural networks. If we are able to extract the brief rules existing behind designs of the complex networks, we will be able to reach better answers on how the various functions are realized within the system. From this perspective, quantitative evaluations of network variables of brain networks have been conducted at various spatial scales.

We see the brain macroscopically, the linked elements through structural networks, white matter fiber bundles, correspond with brain regions. We see the brain macroscopically, the linked elements through structural networks, white matter fiber bundles, correspond with brain regions. The effective number of brain regions consisting of macro-connectome is on the order of 10^2^ −10^3^ (e.g. [Zaleasky, et al., 2010]), if we don’t use voxel-based analyses. Therefore, the number of elements is currently not a serious problem to characterize community architectures and influential nodes in comparison with gene network, protein network, biomedical network, social networks and so on [Radicchi, et al., 2004; Bales, Johnson, 2006].

Recently, however, the research field which regards cells as elements of networks, micro-connectome, is also becoming more active than before. Currently, network variables in neuroscience are calculated directly from the original networks without compressing their dimensions since the number of neurons is also still around 10^3^ −10^4^ in many studies even in cutting edge studies. We need to realize the fact that the simultaneously recordable number of neurons growing twice has doubled each year per 7 years since 1970 [Stevenson, Kording, 2011; Hong, Lieber, 2019]. Of course, the bigger the size of such neuronal networks, the more critical it becomes for us to effectively compress the size, not only to obtain a clear understanding, but also to avoid undesirable properties such as the "curse of higher dimensions”[Bishop, 2006]. Therefore, we needed to prepare a new analysis scheme for any potential technological advances into the future.

### 1-2. How do we compress information?

#### 1-2-1. Network variables

With recent expansion of big data of complex networks including social networks, gene regulatory networks, and real neural networks, various network metrics have been developed to investigate their statistical and topological characteristics of them [Barlebach, Shamir, 2008; Barabasi, Oltval, 2004; Borgatti et al., 2018]. In past network neuroscience studies, various network metrics have also been used to characterize network architectures of individual neural systems [Rubinov, Sporns, 2010; Bassett. Sporns, 2017]. Briefly speaking, as The representative metrics, centralities, clusters, and communities characterize different scale topological architectures of network systems.

First, centralities are metrics to identify important nodes in the network based on various aspects in various (e.g., degree, k-core, page rank, subgraph, etc), and individual nodes are separately provided as relative positions in the entire network patterns [Baveals, 1948; Sabidussi, 1966; Freeman, 1978; Seidman, Stephen, 1983; Brin, Page, 1998; Estrada, Rdriguez-Velazquez, 2005; Borgatti, Everett, 2006].

Second, cluster coefficients and network motifs are metrics relating with small groups of several nodes, and they characterize the statistical frequency of how much individual patterns of several nodes (components) occur than the expected probability expected [Watts, Strogatz, 1998; Milo et al., 2002].

Third, community architectures are global groups of nodes connecting well with each other within the individual groups, which are defined based on various criteria [Girvan, Newman, 2002; Guimera, Amaral, 2005; Lancichinetti, Fortunato, 2009; Kawamoto, 2018]. After dividing networks into several communities, participation coefficient additionally evaluates probability for individual nodes to participate in multiple communities.

Such approaches have the advantage in that it is relatively easy for humans to are relatively easy to interpretthe somehow characteristics of networks of by quantifying individual network metrics. Briefly, these methods provide ability to clear explainability this clearly for humans. This would be because individual variables have been designed based on strong intentionality of individual past researchers that they wanted in wanting to understand specific aspects of the target systems of complex networks.

Although there are excellent virtues of these various evaluation schemes, we will also need to notice that there is no guarantee that the metrics that have been prepared in the past are the ones that are able to optimally characterize the individual datasets that are newly produced. If we see the research fields more widely, extracting features that naturally and optimally match the characteristics of the individual datasets without the researcher’s own subjective bias to these specific network metrics is now becoming a core interest of modern neural network analyses.

#### 1-2-2. Neural network embedding by artificial neural networks

Recently, researchers have used network embedding approaches based on various data-compression techniques to automatically extract features from big complex networks. Among them, "Deep Autoencoders" are a subcategory of deep neural networks that have been historically used and now we have clearly increased the capability with the benefit of recent advances of neural networks. A deep autoencoder is composed of multiple “encoder” and “decoder” layers. The encoder layers transform or compress original data into compressed features’ space, while the following decoder layers recover original data from the compressed features.

Weights of links connecting the nodes in two different layers are optimized through minimizing the inconsistencies between input and output layers. After the process of optimization, the encoder layers with the smallest number of nodes (middle encoder layer) gives a compressed space that captures key features (compressed features) embedded in the original nonlinear complex data [Hinton, Salakhutdinov, 2006]. Because this process enables us to embed the original connectivity data into a compressed features’ space through a neural network, we simply call the approach “neural network embedding”(NNE) in this manuscript.

The weakness of this approach is that it is not easy to provide mathematical interpretation in exchange for the benefits of automation. Therefore, in order to provide the clear meaning for features extracted by the NNE approach, it is necessary to calculate various not only by known but also by newly designed network metrics.

### 1-3. The main target of this study

In this study, we applied one of the training models of artificial neural networks as one of the NNE methods [Wang et al., 2016; Cao et al. al., 2016] to naturally compress the real data of neural connectivity networks into small recoverable data. It is pertinent to note here that there were no cases where NNEs were applied to real neuronal interaction networks.

We demonstrate that our method is able to compress the network architecture, which originally consisted of one hundred neurons in an individual dataset, into 25 features. In other words, the original connection patterns in real neuronal interaction networks, we also call them effective networks, can be recovered from only 25 % compressed space.

Because previous studies demonstrated that network architectures of effective connectivity of neuronal microcircuits includes highly central hubs [Shimono, Beggs, 2014; Schroeter et al., 2015; Nigam et al. 2016; Gal et al, 2017; Kajiwara et al. 2021], this study attempted to observe if this point of attention, that centrality may be a key feature to characterize the network architecture, is still correct in the results of an automatic, non-biased extraction technique. Therefore, we compared the features, compressed in a data-driven manner by the NNE method, with commonly used network metrics, especially various centrality metrics, and several non-centrality metrics. Besides, we also originally designed new network metrics to compare with the features in order to complement the insufficiency of commonly used metrics.

Additionally, we also compared the recovery level of somehow randomized data after passing through NNE, which was trained with original non-randomized data, to check a guaranteed range of recovery from the 25 features.

## 2. Materials and Methods

### 2-1. Physiological recording of neuronal activities

We used neuronal spike data recorded and studied in detail from our past study. So, we briefly explain the experimental procedure utilized in the past study [Kajiwara et al., 2021]. The whole experimental processes are also now open in a video journal [Ide et al., 2019]. We used 7 female C57BL/6J mice (n=7, aged 3-5 weeks). All animal procedures were conducted in accordance with the guidelines of animal experiments of Kyoto University (KU), and have been approved by the KU Animal Committee.

This study recorded neuronal spikes from 300μm-thick cortical slices with a cutting-edge Multi-electrode array (MEA) system (Maxwell Biosystem, MaxOne) with refluxing an artificial cerebrospinal fluid (ACSF) solution that was saturated with 95% O_2_/5% CO_2_ [Kajiwara et al., 2021; Ide et al., 2018]. The slicing positions are accurately controlled with sufficiently less than one slice resolution within somatomotor areas by comparing between 3D scan images of brain surfaces just after extracting brains from heads and MRI images recorded before extracting brains (figure 2a). The morphology of the cerebral cortex, the most recent evolutionary brain region, is like a sheet that envelops all other brain regions. The cortical sheets have a pattern in the depth direction that looks like the overlapping of several sheets, akin to a “baumkuchen”.

**figure 1.**
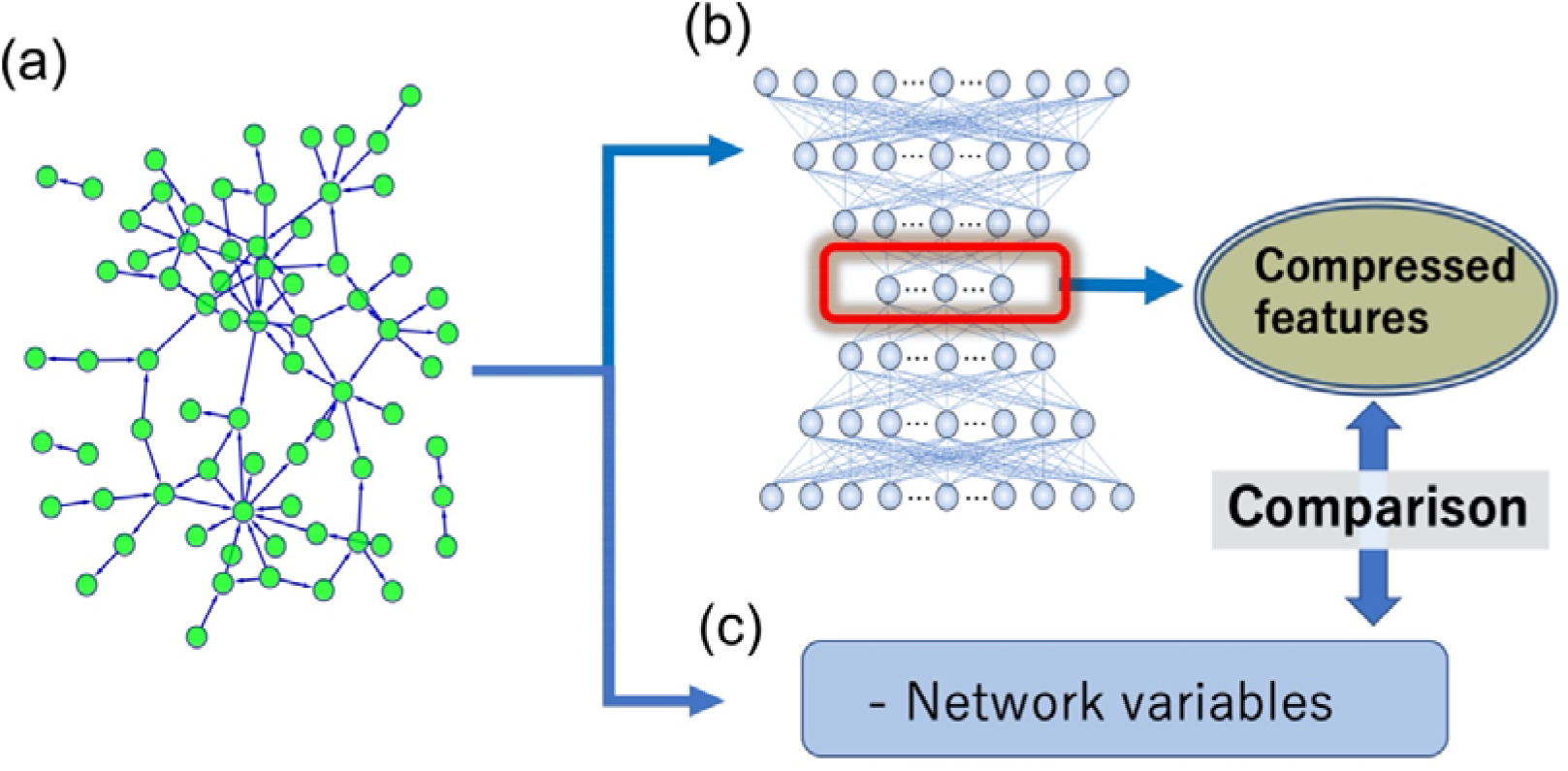
the main logical structure of this study: (a) shows an example of interaction networks among neurons. How we extract compressed features of such networks is the main question that this study asks. (b) The neural network embedding scheme naturally compresses original big networks to a small space through optimizing weights of NNE to the raw connectivity network data. (c) We compared the compressed expression (features) with representative network metrics (e.g., degree, betweenness centrality, subgraph centrality, participation coefficient, clustering coefficient, and new network metrics given in this study). We defined and named the two new network metrics, indirect-adjacent degree and neighboring hub ratio, to improve the explainability of NNE based analysis, and will explain about them later.

**Figure 2.**
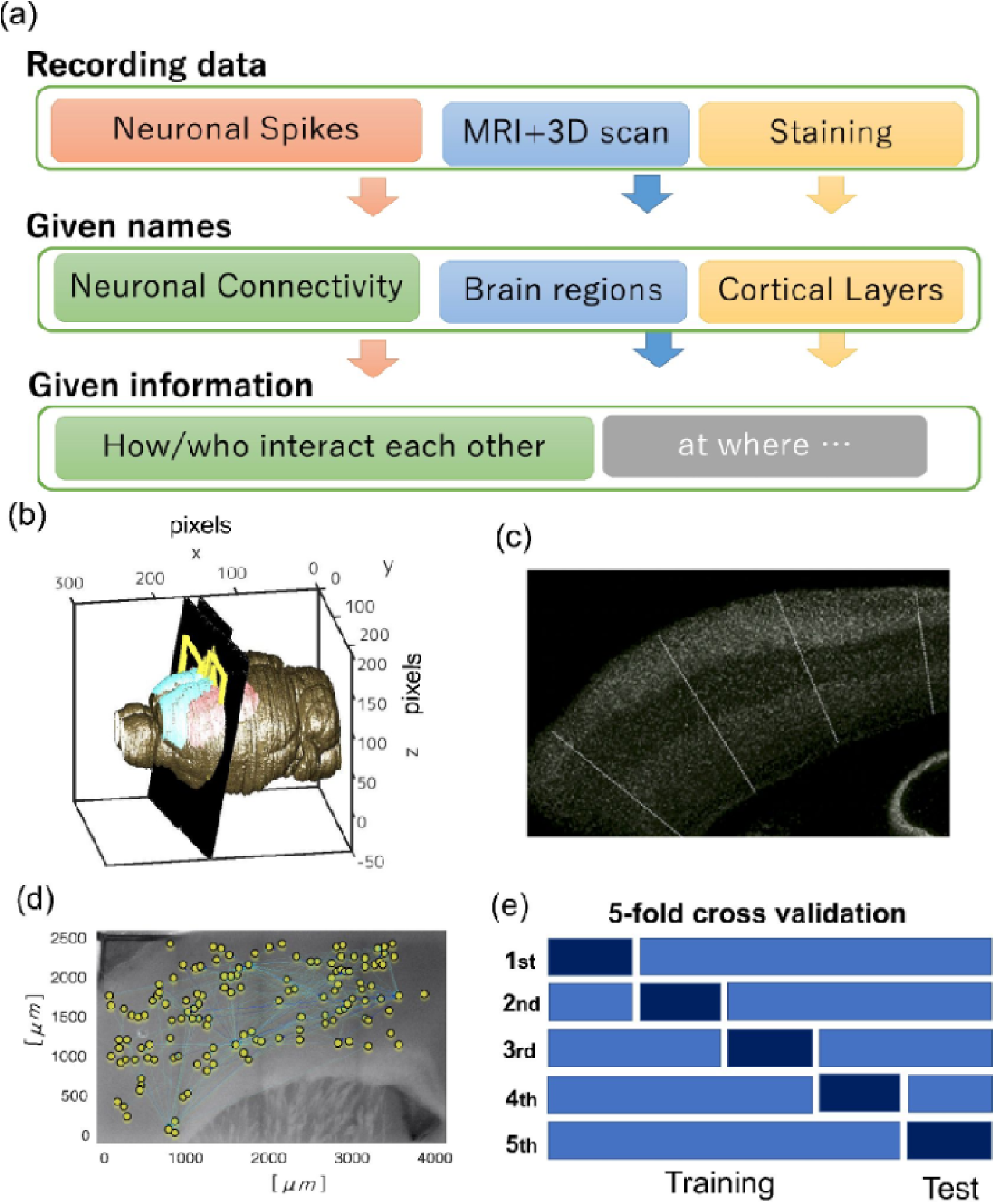
(a) Schematic flow of experimental dataset given from Kajiwara et al. (2021). Read the previous report about the experimental procedure in detail. The given names (categories of physiological targets) and information provided by analyzing the given data are also shown here. We were able to know how (as hub, as making clustering form etc.), which (excitatory or inhibitory neurons) interact with each other at a layer in a brain region. (b) The black slices and yellow regions in the three dimensional brain volumes depict the two dimensional brain surfaces where we recorded electrical activity. (b) This image shows how we are able to know the recording brain region from MRI and slicing position recorded with 3D scan. (c) The white dots are neurons detected with immunostaining, and we are able to find striped patterns in the cortical slice. The pattern is called cortical layers. As we also mentioned in the main document, please notice the meaning of “layer” is different from the number of layers used for defining our artificial neural networks utilized as the analyzing side. As shown as white lines on the cortical slice image, we divided neurons into small subsets holding just 100 neurons, and produced 15 datasets. Because the dividing lines are selected vertically against the layer’s boundaries, we are able to include all layers into all datasets. (d) An example of a connectivity network among neurons plotted on a photo image, in which nodes and links represent neurons and wirings (causal interactions) between pairs of neurons, respectively (see 2-2 for details). (e) Then, we performed 5-fold cross validation with separating the 15 datasets to training and test data. Individual test data included 3 datasets.

We experimentally identified the cortical layers in off-line imaging processes by comparing with NeuN immunostaining images. In our artificial neural network called NNE, the depth of the artificial cells in the input to output direction is also commonly called “layer”. Notice, however, that these cortical layers are a completely different concept.

We also performed spike sorting (Spyking Circus software) to identify ~1000 neurons from the electrical signals, originally, of individual electrodes [Ide et al., 2019; Kajiwara et al., 2021]. The short intervals (15μm) among distances of electrodes for the MEA system enable us to estimate spatial positions of neurons very accurately.

### 2-2. Defining connectivity reflecting causal neuronal interactions

The main target of this study is that we extract and interpret the compressed topological principle representing causal interactions among neurons (figure 2d). Historically, there were many accumulated studies to quantitatively characterize such causal interactions, and past studies of this research field have named networks of causal interactions as *effective networks* [Friston 1994; Aertson, 1989]. Transfer entropy (TE) is one of excellent approaches to estimate *effective networks*, and the topological architecture of reconstructed network using the variable has been studied continuously in various studies by different research groups [Wibral et al., 2013; Lizier et al., 2008; Shimono, Beggs, 2014; Garofalo et al. 2009; Stetter et al. 2012; Orlandi et al. 2013; Kajiwara et al., 2021]. Moreover, TE has shown several preferred abilities for the estimation ability [Lungarella et al., 2007] and also known systematic relations with other information variables [Oizumi, et al., 2016].

This study applies a Network Embedding approach onto effective networks previously estimated from the neuronal spikes [Kajiwara et al., 2021]. Please refer to this past study to know it in more detail.

Before feeding into NNE, we divided the neuron groups into small groups including just 100 neurons, and produced 15 datasets. Then, we divided neuron groups by a dividing line in vertical to the cortical surfaces (white lines in the figure 2c). Because of these divisions, all datasets include all cortical layers and each neuron can participate in only one data group. We sorted the order of neurons according to ([cell category] - 0.5)*[layer category] to reasonably express neurons’ identities with one index. Here, the [cell category] is 0 for inhibitory neurons and is 1 for excitatory neurons. The value [layer category] is one of four indices 1,2,3,4 expressing cortical layers 1-3, 4, 5, 6 respectively.

We separated the 15 datasets into 5 sub-datasets and used one of 5 sub-datasets as test data, and remaining 4 sub-datasets as training data in five hold validation procedures (figure 2e). Neurons (nodes) having no links about both inputs and outputs were eliminated in the following procedure because we can estimate such neurons are effectively isolated in the slice.

### 2-3. Network embedding: Searching minimum expressions

#### 2-3-1. Neural Network embedding (NNE): non-linear automated feature extraction

In performing neural network embedding (NNE) analysis, in order to simply characterize network architectures, we used an artificial neural network having a symmetric layer architecture composed of multiple encoders and decoder layers. Specifically, the number of units in each layer, so at each depth of the artificial neural networks, the number of artificial neurons(units) decreases linearly from the number of experimentally recorded neurons given in the input layer. Then, the number increases linearly again from the middle layer to the output layer. This allows the design of the NNE to be regulated by only two parameters, the depth of layers and the number of middle layer nodes (see Figure 3-a). These layers are fully connected and those except for the output layer use the activation function of the rectified linear unit (ReLU) [Brownlee, 2019]. The output layer used activation sigmoid function to transform output values into binary values.

**Figure 3.**
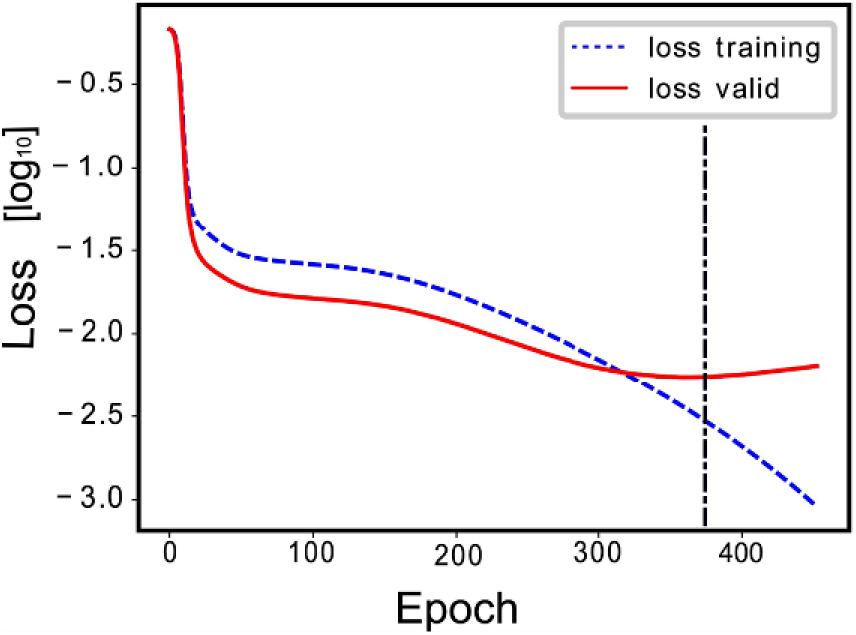
A schematic example of learning curves for training data and for validation data. The figure shows how the value of loss changes as the number of epochs increases and the NNE model is trained. The dotted line shows a result for the training data, and the solid line shows a result when the trained model is applied to the remaiing test data.

**figure4.**
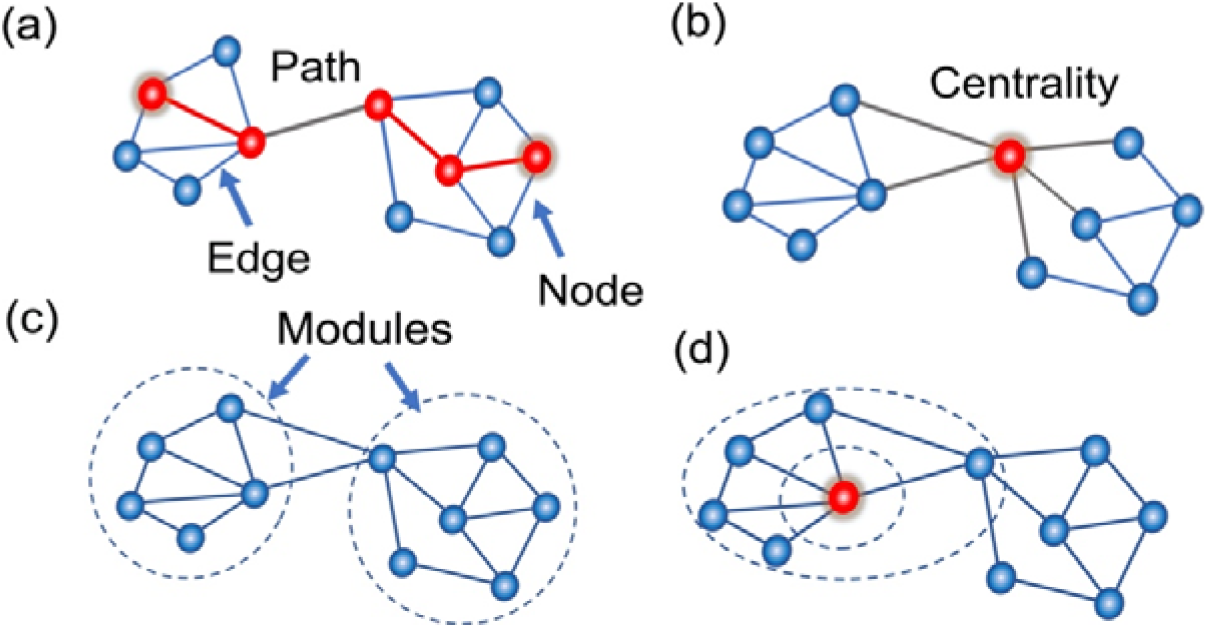
Concept figure to explain utilized network metrics: This panel (a) illustrates the concepts of nodes, edges, and paths. A node is a point connected by an edge (line), and a path is a series of paths to go from one node to another node connected indirectly. Panel (b) illustrates centrality. It evaluates how central a node is in the whole network from various perspectives. For example, degree centrality is evaluated based on how many nodes are connected to it, while betweenness centrality is evaluated based on how many of the shortest paths between nodes pass through it. (c) A module is a large group of nodes in the overall network. The density of connections within each module is high, while the density of connections between different modules is relatively low. (d) The ability given in common to the newly defined network metrics is to find situations where hubs are located at some path length away from the particular node of interest (red dot), or where an increase in degree can be observed.

We optimized the artificial neural networks based on the “Adam” optimizer [Kingma, Ba, 2015], and selected default values for parameters of the optimizer (the default values of learning rate, beta 1, beta 2, and epsilon, see manual document for “Adam” function in Keras, _https://keras.io/api/optimizers/adam/, for details). In the optimization step, discrepancies between values in the input layer and those in the output layer, quantified with binary cross-entropy loss, were minimized as the common scheme of NNE optimization.

This study divided the 15 datasets into 5 sub-datasets and selected one of the five as a test dataset. The remaining four sub-datasets were used for training dataset. Then, we trained the NNE model with five different combinations of datasets and evaluated the loss function as the average value. When the loss function did not decrease for 50 consecutive epochs in the training data, we regarded that the 50 epochs as sufficient to converge, and used the last epoch count for the final evaluation of the test data. We implemented all NNE models using a deep learning platform Keras [François Chollet. Keras. https://github.com/fchollet/keras, 2015] with Tensorflow (www.tensorflow.org) backend.

#### 2-3-2. Evaluation of reconstruction errors

Reconstruction errors evaluate losses (or inverse of performances) measuring discrepancies between inputs and outputs after passing through a compressed latent space by embedding methods including NNE, i.e., lower reconstruction errors for embedding methods indicate that their embedding performances are higher. The training process of the NNE model runs toward minimizing the errors. Here, the reconstruction error was quantified with binary cross entropy loss to optimize the NNE models as we briefly mentioned in the previous subsection. The binary cross entropy loss is well-known as a measure that is more compatible with stochastic gradient descent than mean squared error [Goodfellow, Bengio, Courville, 2016], and is defined by the following equation:

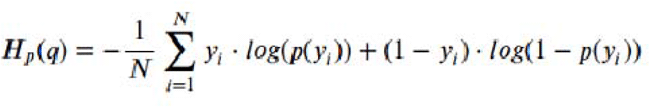

We also utilized the binary cross entropy loss to quantify how much the reconstructed signal from swapped data through trained NNE models differs from the reconstructed one trained from raw data.

### 2-4. Network metrics: Interpretations of extracted features

#### 2-4-1. centrality metrics and several other network metrics

In order to give interpretations to the automatically extracted features by NNE analyses, we calculated 13 network metrics because their quantitative meaning is very clear for researchers. Among the calculated network metrics, the main first group of metrics is centrality-type metrics such as degree, subgraph centrality, betweenness, closeness, page rank. The centrality-type metrics are focusly selected selected according to focus because past studies found that centrality or the closely related measures are a key feature characterizing the network topology of local effective neuronal connectivity networks [Shimono, Beggs, 2014; Nigam et al., 2016].

Here, let us explain the basic definitions of those centrality metrics. As the most basic centrality metric, degree of node i is the number of nodes that have a link to the node i. Betweenness defines centrality in terms of paths, connecting nodes between two different nodes. For example, betweenness for a given node is the number of shortest paths connecting all possible pairs of nodes that pass through the node i. Alternatively, Subgraph centrality quantifies centrality in terms of small subsets of nodes, subgraphs. Subgraph centrality of a node i is the ratio of how much the node often/widely participates in many different subgraphs included in a network [Estrada E and Rodriguez-Velazquez JA 2005].

In addition, the Cluster coefficient of a node i (Ci) is calculated by using the following equation. Ci = 2ei/ki(ki-1), where ki is the degree of the node i and ei is the number of links connecting neighbour nodes of the node i to one another [Watts and Strogatz, 1998].

In order to compare these centrality metrics, we also calculated participation coefficient which is a metric defined based on community structure. Although participation coefficient is relatively close to subgraph centrality, the participation coefficient defines how often a node participates in many various communities. The values represent the big architectures of networks because the basic group nodes are not small subgraphs but big communities.

We used ‘NetworkX [Aric A et al 2008]’, ,’python-louvain’, and ‘Brain Connectivity Toolbox [Rubinov, Sporns, 2010]’ python modules to calculate these representative network variables.

#### 2-4-2. designing new metrics

Since we could not sufficiently give interpretations for several compressed features of NNE with the centrality metrics, we were obliged to design new network metrics. Therefore, we attempted to design new network metrics having supplemental properties to centrality measures. We have two network metrics that are expected to account for topological features that could not be covered by centrality or other common network metrics:

We named the first new network metrics n-step neighboring hub ratio; The metric quantifies the ratio of hub nodes among nodes n-step apart from the node i. Here, the hub nodes were defined as nodes having the top 20 percent highest degree [Yu H et al 2007;Barabasi AL 2016]. We examined cases of n = 1, 2, 3, and 4 for all the nodes in the neural connectivity maps by considering the matrix size. These metrics are inspired by several past studies discussing the “second neighborhood problem” [Chen et al., 2003; Brantner et al., 2009].

Besides, in order to capture and characterize more efficiently, we prepared another type of new metric. The second new network metric is named relative Indirect-adjacent degree, which assigns a high score for a node two-steps apart from hub nodes. More specifically, this metric is calculated by using the following equation <D_2_(i)>/<D_1_(i)>, where <D_1_(i)> is mean value of degrees among neighbor nodes of a given node i and <D_2_(i)> is mean value of degrees among nodes two-steps apart from the node i. We could expect this variable to also emphasize nodes slightly far from hubs. and which may emphasize around boundaries rather than around central modules. This variable succeeded to capture one of NNE features uncorrelated with centrarities. Refer figure 7 (c),(d), figure 8 (e) to observe their distributions more directly.

### 2-5 mutual information

We used mutual information to investigate relationships between NNE’s compressed features and 13 representative network metrics [Studholme C et al 1998]. We used codes modified from https://github.com/mutualinfo/mutual_info to calculate mutual information.

## 3. Results

### 3-1. Compressing with Neural Network Embedding

In order to optimize the architecture of Deep Neural Networks to achieve high reconstruction performance of input network architecture, we attempted to minimize the binary cross entropy loss that characterizes gaps between input networks and their reconstructed output networks. The input networks are the connectivity matrix representing causal interaction among neurons [Kajiwara et al., 2021].

Here, the number of units, artificial neurons, at individual layers were once *linearly decreased* from the number of experimentally recorded neurons given on the input layer to the minimum number of neurons located at the most middle layer (Fig. 5-a). Following the decreasing, or compressing phase, the number of units is linearly increased again from the most middle layer to the output layer. Of course, the number of units in the output layer is the same as the number of units in the input layer. Because of the network design, the architecture of the deep neural networks could be characterized only with the pair of parameters, *depth of layers (Depth)* and *number of neurons at the middle layer (Middle Size)* (Fig. 5-a).

**Figure 5.**
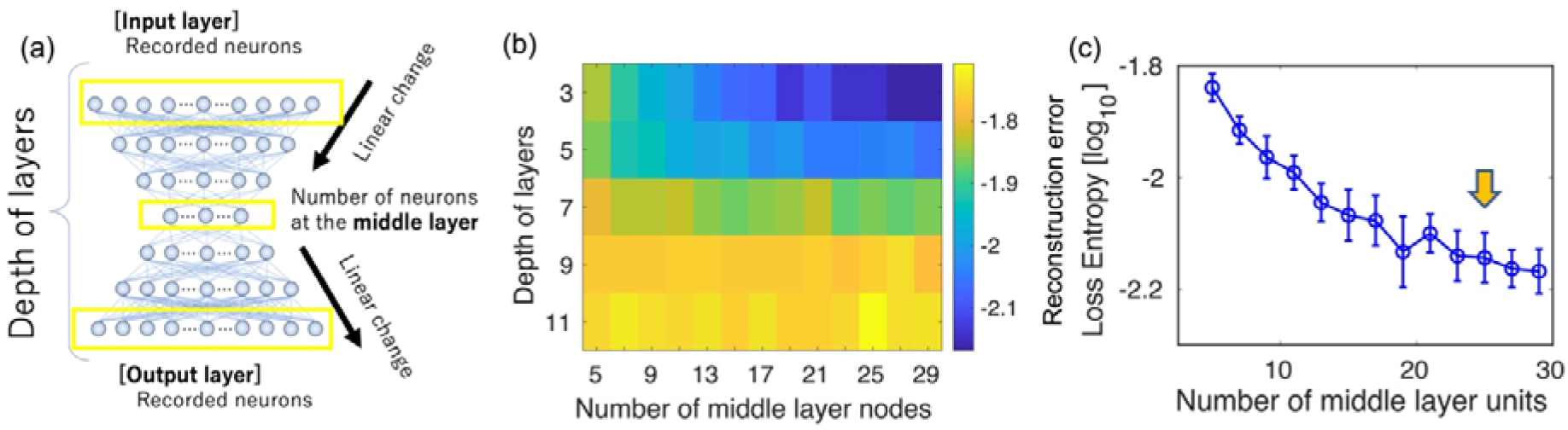
the network architecture of deep neural networks and loss in the learning process. (a) The architecture of neural networks utilized for deep neural networks in this study. The number of units (artificial neurons) gradually decreases in a linear trend from the number of units of the real neural network to a much fewer number of units located at the middle layer, and gradually increases in a linear trend again toward the output layer. The key two parameters describing this network architecture are (1) the depth of layers, and (2) the number of units at the middle layer. (b) The loss function, binary cross entropy, is mapped on the two dimensional map of “depth of the layers” (“Depth”) and “number of middle layer units” (“Middle Size”). The binary entropy was averaged for seven data samples. (c) Binary cross entropy only on the section at depth layers equal 3 is shown. We especially focus on the parameter region where the “number of middle layer units” is 25 shown as the inserted allow.

Based on the basic architecture we observed loss functions for “Depth” between 3-11, and “Middle Size” between 5-29 (fig. 5-b). We performed five fold cross validation after separating training data (12/15) and test data (3/15). Refer the detailed learning and test processes to the method section.

After sufficient learning steps, which were adaptively chosen to reach a plateau of loss function, the cross entropy both for validation data. From the colormap of loss function for test data (fig. 5-b), we could find the loss gradually increases when we make the network architecture deeper. This might be caused by gradient disappearance even though we used ReLU function, which has a good ability to avoid the phenomenon.

The result of the initial survey of wide-range parameters nicely suggested that Deep Neural Networks is able to compress the size of complex networks into 25% nodes or units while stably keeping the loss less than 10^(−2.2)) for all data sets. Based on the wide survey, we selected the pair of parameters (Depth, Middle Size) = (3,25) as the representative one because of the following reasons: The first parameter 3 of Depth was regarded as the optimal size about the depth of layers because the cross entropy loss is lower than other deeper (>5) networks. The second parameter was selected because the values of the loss on the section of the depth of layers equals 3 once reached a plateau value around where middle size is 25 (pointed with an arrow in Fig. 5-c).

### 3-2. Interpreting compressed features with centrality measures, and other common network variables

Here, we calculated five centrality network variables: degree, subgraph centrality, betweenness centrality, core number, and pagerank. Unfortunately, none of these variables showed a particularly more significant correlation with broad NNE features than the others. In the end, these centrality variables were likely to be able to interpret ~30% of the features of NNE (Figure 7-e).

In order to intuitively observe the features that are relatively similar to the centrality metric, we drew a network in which the 8th features from the NNE model are reflected in the sizes of the markers (Figure 6-c). As shown these results, we can see that the larger markers are naturally placed in the center of the network.

**Figure 6.**
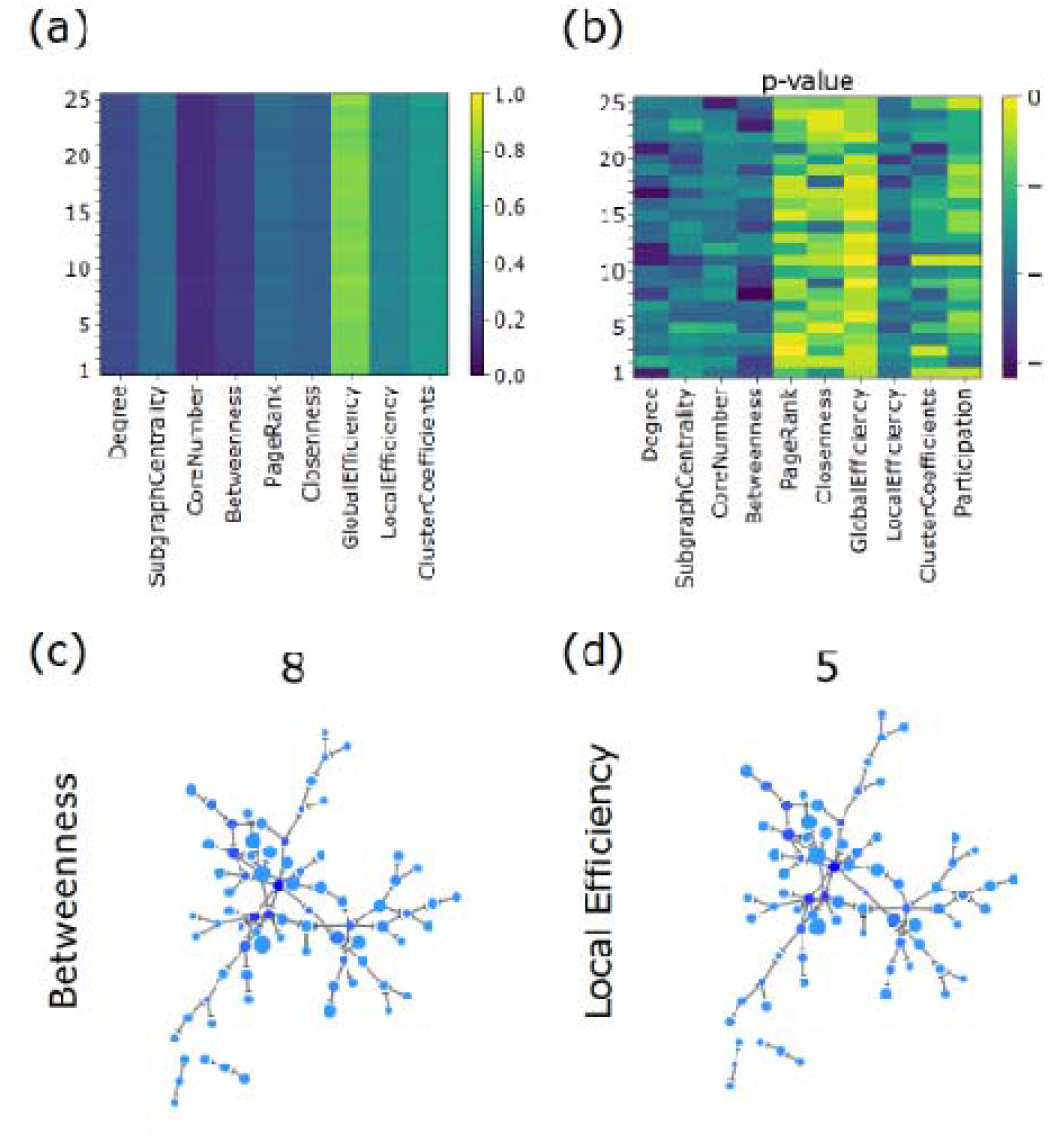
Comparing NNE’s features in comparisons with Network variables: (a) This colormap shows averaged mutual information between NNE’s features and several commonly utilized network variables, including the 6 centrality-type and other type metrics. (b) is the result of a two-sided Wilcoxon’s signed rank test comparing from zero point. The blue regions express pairs of variables showing relatively smaller p-values, and we regarded it as significant if the p value is smaller than 0.005. (c) is an example of network visualization with expressing betweenness centrality as marker sizes and 8th NNE’s feature as the marker colors. This graph visually represents the tendency for nodes with high centrality to have strong values for the 8th NNE’s feature. (d) is an other example between local efficiency and 5th NNE’s feature.

In addition to centrality variables, we also prepared different types of common network variables, such as local efficiency and participation coefficient. We, however, could not characterize ~40% NNE features with any common network variables (figure 6-d, 7-e).

**Figure 7.**
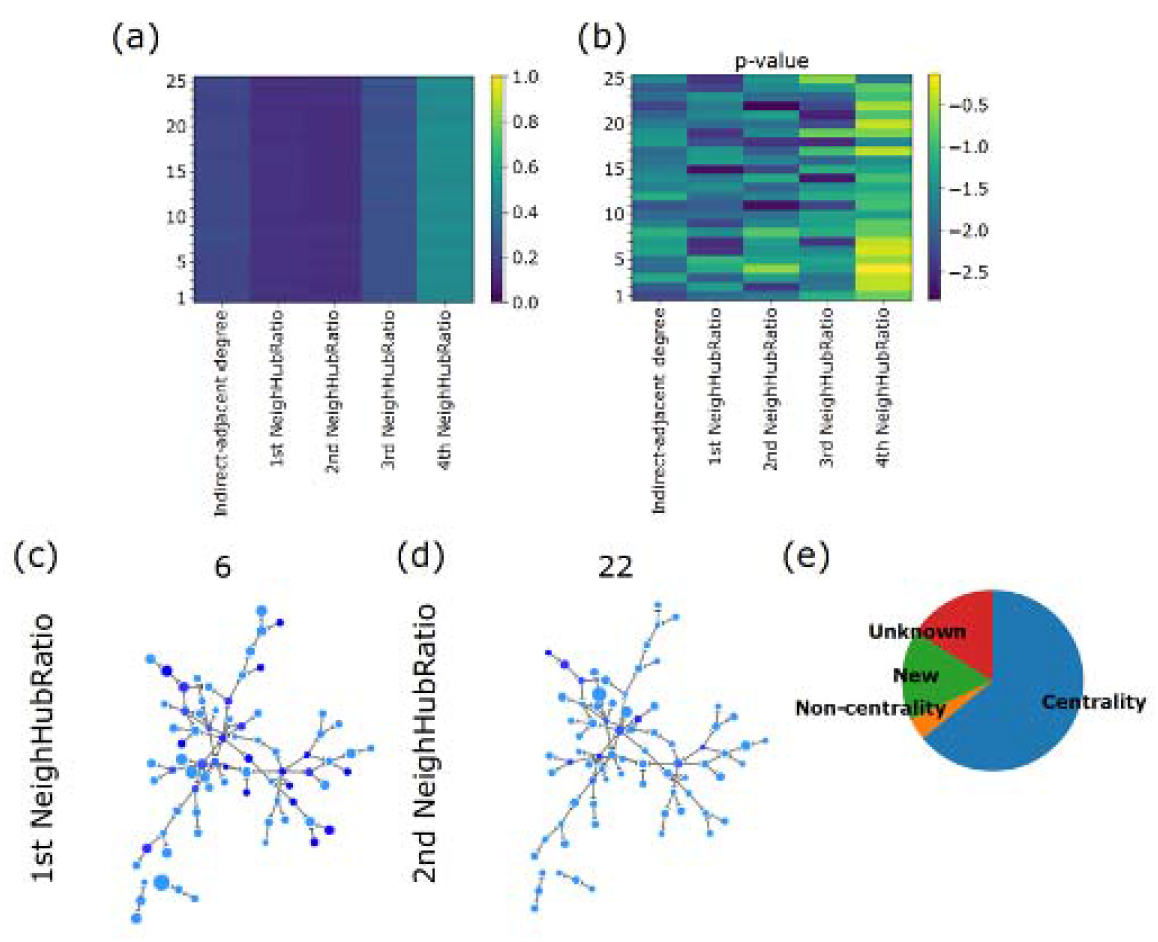
Evaluation of NNE’s features with newly designed network metrics: (a) The color map of mutual information between NNE’s features and newly designed network metrics, indirect-adjacent degree and several order Neighbour Hub Ratios (Refer the method section about their definitions). (b) The result of Wilcoxon’s rank sum test compared with zero points. The significant correlation (p<0.001) is colored as yellow. (c),(d) Example of network visualizations to show relationship between NNE features and representative network metrics. Marker sizes reflect network metrics and marker colors show NNE’s features written in the individual panels. (e) The ratio of Centrality type, Non-centrality type and Newly designed network metrics significantly relating to NNE features. The new metric played a fairly good role to interpret the automatically extracted features.

### 3-3. Explanation ability by adding originally designed metrics

Can we selectively interpret the characteristics of the compressed features from the NNE models that did not relate to commonly used network variables? - In order to capture such characteristics, we proposed two new network metrics (Figure 7-a, b). The first metric that we designed is Indirect-adjacent degree, which is ratios of between degrees of neighbor nodes of a given node *i* and degrees of nodes two-steps apart from the node i. The second metric that we designed is N-st Neighboring Hub Ratio, which quantifies the ratio of hub nodes, nodes with top 20 percent highest degree, among nodes N-steps away from a given node *i* [Yu H et al. 2006, Barabashi AL 2016].

Although we could expect that these metrics clearly have the ability to capture characteristics that did not significantly relate to common centrality and non-centrality type network metrics, the variable succeeded to capture many features more than expected. The ratio of newly captured features was around 20% (figure 7-e).

Now, let us draw two networks in which the 6th and 22nd NNE’s features are reflected in the marker size (figure 7-c,d). These examples demonstrate that the maximum marker size was reached at one or two nodes away from hubs. This property is consistent with our new metrics, neighbor hub ratios.

Besides, there were also still several unknown features, as shown in the figure 7-e. So, more future effort will be necessary to answer the open question: what network variable will capture such non-uniformity of such still “unknown” features.

### 3-4. Transposed case

Next, we used the transposed matrices, where the rows and columns of the matrices were exchanged, as input to the NNE. This analysis implies that for a particular neuron, instead of analyzing the input signals, it analyzes the output signals. As a result, we observed basically the similar trends that were observed in the non-transpose case (fig.8).

**Figure 8.**
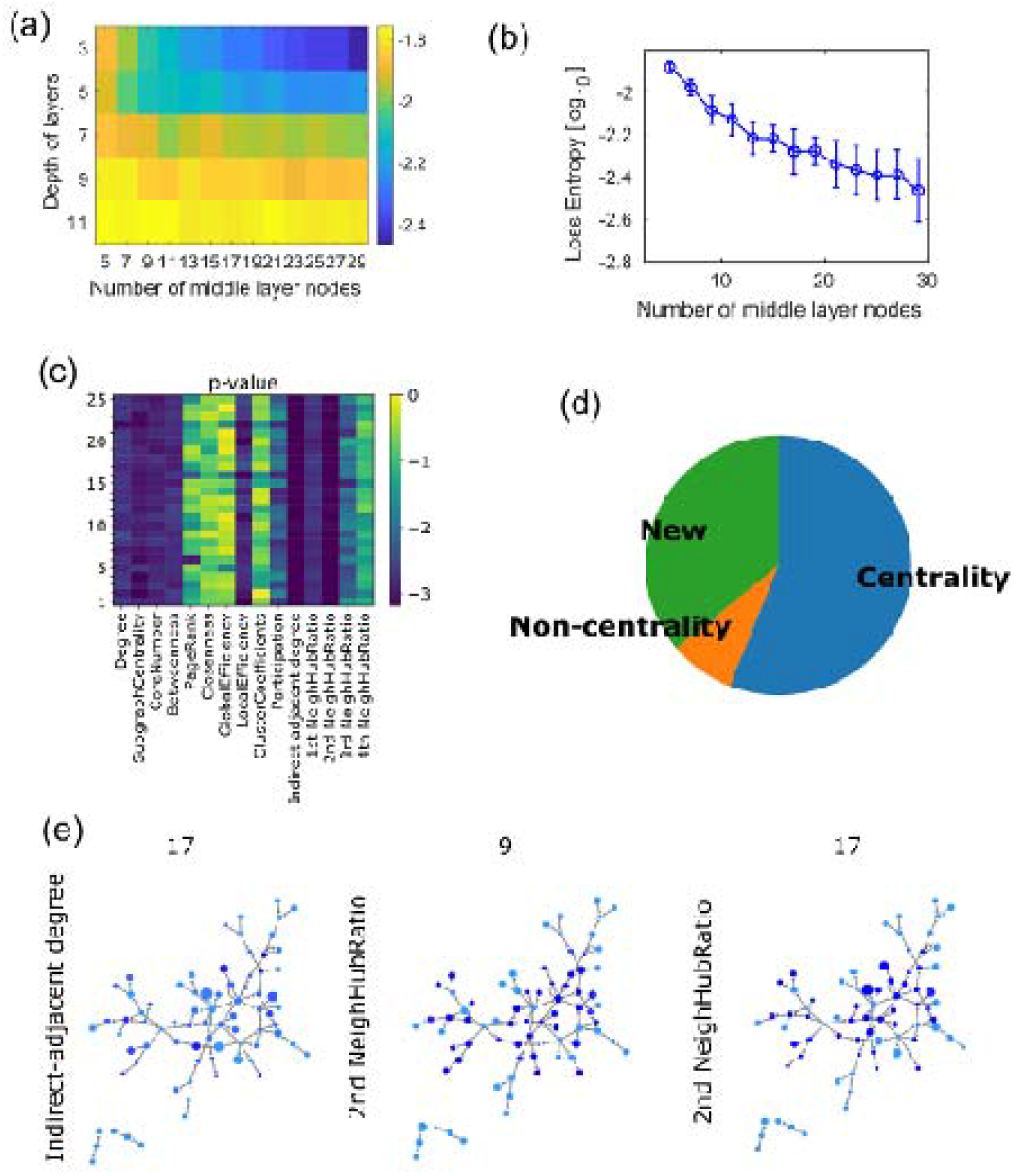
Evaluations of transposed matrices: (a) The influences of two parameters, Number of middle layer nodes and Depth of layers, of NNE’s architecture. This panel is the same map with figures 5-(b) except that we utilized the transpose matrix as the input to the NNE. (b) is also basically the same as figures 5-(c) except the input adjacency matrices were transposed. (c) is the colormap of p-values for Wilcoxon’s ranked sum test applied to mutual informations between NNE’s features and originally designed network variables here (d) To sum it all up, the relative contribution of common centrality metrics, non-centrality metrics, and newly designed metrics are shown as the pie chart here. (e) shows three examples of network visualizations, in which marker sizes express network metrics and marker colors express NNE’s features written in the individual panels at the positions of y-axis and title.

The most important point among the common characteristics is that the network metrics commonly used in the past, mainly centrality type metrics, significantly relate to around 50% percentage of the 25 compressed features. Surprisingly, as much as ~100% of the metrics could be given interpretation with the addition of the new metrics (fig. 8-d).

In particular, in the case shown here, the Indirect Adjacent Degree and 2nd Neighborhood Hub Ratio are significantly related with a wide range of features, and are expected to capture the important characteristics extracted by NNE (fig.8-c, d, e). In summary, we seem to be able to say not only there is a tendency about relation with network metrics similar to non-transposed case, but also that there is also a tendency for loss to fall further when transposition is performed compared to when no transposition is performed.

### 3-5. Evaluation in swapped networks

It is still unclear from the analysis so far how much generalization ability the NNE model has or how much specialized information is extracted from the real data. Therefore, in order to evaluate them quantitatively, we examined whether the error could become larger when we input swapped data to the NNE model, which accomplished training with real data.

The swapped data was generated by randomly moving the connected part of the original connection matrix to another place in the matrix while preserving the out degree histogram [Maslov m, Kim, 2002]. The percentage of the number of swapped connections (relative to the original number of connections) was gradually increased to 0, 20, 40, 60, 80, and 100 % [fig. 9-b]. The results showed that information loss occurred and the Cross Entropy measuring the loss gradually increased. The increase of Cross Entropy began to occur at the point of 20% swapping, which suggests that the NNE learns specific patterns on the raw data without swapping.

**Figure 9.**
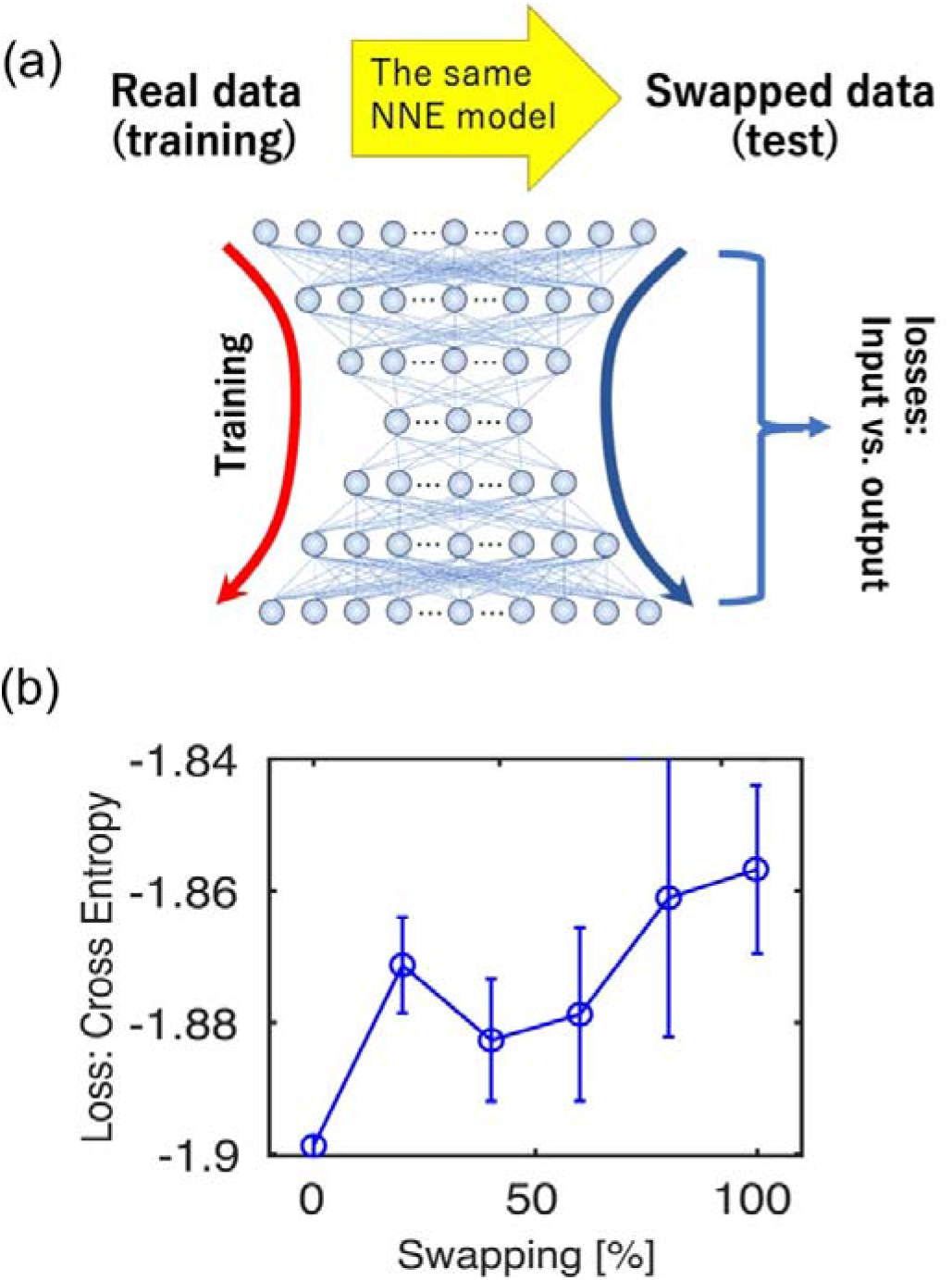
the evaluation of generalization ability to swapped data. (a) As shown in the left panel, we first trained the NNE model with read data. After that, we evaluated how much the error increases when the swapped networks are fed to the trained NNE (right panel). (b) When the degree of swapping to the network was gradually increased as a percentage of the number of connections that originally existed, we evaluated how much error (information loss) the restored network had using Cross Entropy after compression through NNE. The errorbar is standard deviation for 5 time repetitions.

## 4. Discussion

In interpreting data, it is crucial to eliminate as much of the researcher’s bias as possible because final conclusions might be greatly influenced by the metrics that, for example, network scientific metrics, selected at the starting point of the research project. As mentioned in the introduction, the number of neurons that can be measured simultaneously on the nervous system is exponentially increasing, and the development of automatic compression technology is expected to become gradually more important in order to compress the large information in the near future.

From these backgrounds, this study proposes an “neural network embedding” (NNE) that automatically extracts features from the network architecture without the subjective bias of the researcher, by utilizing artificial neural networks, which have undergone considerable development in recent years.

This research aimed to provide an interpretation of the features extracted by the NNE in a data-driven manner by comparing them step by step with network metrics.

Firstly, we compared automatically extracted features with centrality metrics. These centrality indices significantly relate to 50-60% of the total features, which means that the importance of the centrality indices is guaranteed even by a bias-free method.

Secondly, we added some other common metrics of non-centrality as the comparing targets. However, the explanatory power of these metrics did not seem to be high. For example, the participation coefficient is a representative network metric based on the module [Rives, Galitski, 2003; Han et al., 2004; Shimono et al., 2014]. The reason why this metric did not show high performance is that we limited the number of neurons to 100 in order to prepare 15 datasets that allow us to evaluate the generalization performance of the learning process. This number of neurons distribute in less than 1/3 area of the measurable surface by our recording system (2*4mm). The typical spatial size of the columnar structures corresponding to physiological neuronal modules of neural local circuits, however, is slightly larger than the spatial area covering ~100 neurons [Shimono, Beggs, 2014].

Finally, we prepared our own network metrics, which we named Indirect-Adjacent Degree and Neighbor Hub Ratio. Both of these metrics focus on situations where neurons that are a few steps away from the neuron (node) of interest correspond with hubs, or have a higher degree than the surrounding adjacent neuron.

These new metrics showed significant relationships with NNE’s features that were not found to be significantly related to the commonly used network metrics, and were thought to contribute to a certain interpretation of these unknown features. This trend was also observed when the network was transposed, but this is not necessarily a trivial result.

Whatever, these results successfully demonstrated that this NNE approach is also expected not only to compress effectively but also to provide opportunities for human researchers to develop new network metrics.

As mentioned in the introduction, this NNE approach will become more important in the future as the simultaneously recordable number of cells increases. Similar network embedding methods are beginning to be applied to the brain-wide connectome, which consists of networks connecting brain regions [Rosenthal et al., 2018]. When considering networks of neurons and other cells as elements, the number of required elements will potentially increase by another 3-5 orders of magnitude. Therefore, such compression methods are more critical.

So far, researchers have developed various network embedding methods based on different strategies, including especially deep learning methods. For example, Perozzi et al. developed a network embedding method, “DeepWalk”, that used random walk of short steps together with SkipGram modeling to capture topological structures of neighborhoodness for each node [Perozzi et al., 2014]. Following it, Grover and Leskovec proposed “node2vec” that improves DeepWalk by using efficient node sampling techniques to capture whole spectrums of connectivity patterns in complex networks.

Recently, wider researchers have developed new embedding methods based on deep learning. Several researchers have proposed deep autoencoder based methods (e.g., SDNE, SDAE, and SiNE) to extract compressed features from a big data of complex networks [Vincent et al 2010, Xu et al. 2021]. Such deep learning based methods show significantly higher expressive powers than other embedding methods. Researchers have leveraged compressed features by deep neural networks for community detection [Ye et al., 2018] and nodes clustering [Yang, et al., 2019] in complex networks. Recently, Tsuji et al. used a deep autoencoder to extract compressed features from a protein interaction network and leveraged the features for inference of potential novel drug-target genes for Alzheimer’s disease [Tsuji et al 2021].

In the future study, such various methods will provide deeper insight or higher predictive power of brain states, including various disease states.

**Table 1.** Subcategories of network metrics: Network metrics are categorized into three types. The first type is the commonly used metric of centrality, which has been widely examined because past studies have shown that nodes with exceedingly high centrality exist in local circuits of the nervous system. The second type is a commonly used metric that is somewhat different from centrality. Although local efficiency is similar to betweenness centrality, it evaluates the ability of individual nodes to shorten paths in a more local network structure (cluster). The participation coefficient evaluates whether a node is able to generate information flow across modules in the whole network (fig.4-c). The third type is a new set of network metrics, which were originally defined in this study to compensate for the characteristics that the centrality metrics might overlook. Refer to the main manuscript for their explanation.

|  |  |  |
| --- | --- | --- |
| **Type 1** | Centrality | Degree , SubgraphCentrality, CoreNumber<br>Betweenness, PageRank , Closeness |
| **Type 2** | Non-centrality | LocalEfficiency , Participation coefficient<br>Cluster coefficients |
| **Type 3** | New Metrics | Indirect-adjacent degree , 1st neighbor hub ratio<br>2nd neighbor hub ratio<br>3rd neighbor hub ratio , 4th neighbor hub ratio |

## Acknowledgement

MS is supported by several MEXT fundings (17K19456, 19H05215, 20H04257) and Leading Initiative for Excellent Young Researchers (LEADER) program, and grants from the Uehara Memorial Foundation. The MRI experiments of this work were performed in the Division for Small Animal MRI, Medical Research Support Center, Graduate School of Medicine, Kyoto University, Japan. We warmly acknowledge Takuma Toba, Tatsuya Tanaka, Hirohiko Imai and all support by Hakubi center to establish this study.

